# KaryoTap-based scDNA seq assessment of aneuploidy induction by hydroquinone in hematopoietic stem cells

**DOI:** 10.64898/2026.11.02.763824

**Authors:** Maria Trifas, Stephanie Aronson, Fabio Alfieri, Manjunatha Kogenaru, Rachel Kim, Teresa Davoli

**Affiliations:** Institute for Systems Genetics and Department of Biochemistry and Molecular Pharmacology, NYU Grossman School of Medicine, New York, NY, USA; Department of Pediatrics, NYU Grossman School of Medicine, New York, NY, USA

**Author notes:** Lead Contact and Correspondence: Teresa Davoli (,).

## Abstract

Monosomy 7 is a frequent cytogenetic alteration associated with poor outcomes in myelodysplastic syndrome and acute myeloid leukemia. The benzene metabolite hydroquinone (HQ) has been reported to induce chromosome 7 loss in cultured hematopoietic cells, suggesting a mechanism linking benzene exposure to these diseases. However, these findings were largely based on FISH analysis of selected chromosomes in relatively small numbers of cells. Here, we tested whether HQ increases aneuploidy in primary human CD34+ hematopoietic stem and progenitor cells (HSPCs) using KaryoTap, a single cell DNA sequencing approach that measures copy number across all 22 autosomes, together with interphase FISH. We first identified 20 µM as the highest HQ concentration tested that preserved sufficient viable cells for single cell analysis. We then examined cord blood and adult mobilized CD34+ cells, including the MPS1 inhibitor reversine as a positive control in the cord blood experiment. Reversine increased whole chromosome gains and losses, including those of chromosome 7. HQ did not produce a reproducible increase in aneuploidy across three donors and two sources of CD34+ cells, including two concentrations and two timepoints in the adult donor. A modest increase in chromosome 7 loss in one cord blood donor was not reproduced in the other donors. In the other cord blood donor, the estimated upper bound on the HQ associated increase was approximately 1.4 percentage points. HSPCs also showed a greater increase in chromosome losses than epithelial cells or melanocytes following the same reversine exposure. These findings do not support a substantial, reproducible increase in whole chromosome aneuploidy after HQ exposure under the conditions tested. They are consistent with a model in which benzene exposure favors the expansion of preexisting monosomy 7 clones, although this possibility requires direct testing.

**Highlights:** KaryoTap measured copy number across all 22 autosomes in primary human CD34+ HSPCs exposed to hydroquinone.

Hydroquinone did not reproducibly increase whole chromosome aneuploidy or monosomy 7 in cord blood or adult mobilized CD34+ cells.

Reversine increased chromosome gains and losses in the same cord blood cultures.

The findings support further testing of whether hydroquinone favors the expansion of preexisting monosomy 7 clones.

## Introduction

Aneuploidy, defined as an abnormal number of whole chromosomes or chromosome arms, is relatively uncommon in healthy tissues but frequent in human cancers, including hematopoietic malignancies (Knouse et al., 2014; Ben-David and Amon, 2020). Loss of chromosome 7 (monosomy 7) or its long arm (del(7q)) is among the most recurrent somatic copy number alterations in myeloid disease. These alterations occur in 5 to 10% of adult sporadic myelodysplastic syndrome (MDS) and in 40 to 50% of pediatric MDS, often as the only detectable cytogenetic abnormality (Kardos et al., 2003; Inaba et al., 2018; Wlodarski et al., 2018; Jotte and McNerney, 2022). Monosomy 7 is associated with clonal progression, transformation to secondary acute myeloid leukemia (AML), and adverse clinical outcomes (Greenberg et al., 2012; Papaemmanuil et al., 2016). Most studies have focused on why monosomy 7 cells expand after the lesion has arisen. Less is known about how hematopoietic stem and progenitor cells (HSPCs) initially lose chromosome 7, or whether associated environmental exposures promote the generation of the lesion, the expansion of cells that already carry it, or both.

Benzene is an established human leukemogen linked to AML and, in pooled analyses of petroleum worker cohorts, to MDS (Schnatter et al., 2012; Shallis et al., 2021). Benzene exposure has also been associated with childhood leukemia, which is relevant given the high frequency of monosomy 7 in pediatric myeloid disease (Reynoso-Noveron et al., 2024).

Benzene has been proposed to promote monosomy 7 both by inducing chromosome loss and by selecting cells that already carry this alteration. Its toxicity is mediated in part by metabolites, including phenol and hydroquinone (HQ). HQ accumulates in the bone marrow, where myeloperoxidase converts it to 1,4-benzoquinone, a reactive electrophile to which CD34+ progenitors are particularly sensitive (Smith et al., 2000; Abernethy et al., 2004).

Studies of benzene exposed workers in China reported dose dependent increases in monosomy 5, monosomy 7, del(5q), and del(7q) in peripheral blood lymphocytes analyzed by FISH (Zhang et al., 1998a). A subsequent study found increased monosomy 7 and monosomy 8 in granulocyte macrophage colony forming cells from workers exposed to benzene concentrations below 10 ppm (Zhang et al., 2012). These changes are not restricted to chromosome 7. An eight color FISH study examining all 24 chromosomes in a separate worker cohort identified dose dependent increases in monosomy of chromosomes 5, 6, 7, 10, 16, and 19, together with gains of several chromosomes (Zhang et al., 2011). Whether benzene associated myeloid neoplasia has a distinctive cytogenetic profile also remains debated.

Analyses of karyotypes and chromosome breakpoints have suggested that benzene associated AML may more closely resemble de novo AML than therapy related disease (Irons et al., 2013; Kerzic and Irons, 2017).

Several in vitro studies have reported that HQ is sufficient to induce chromosome alterations. At concentrations of 25 to 50 µM, HQ increased monosomy 5 and monosomy 7 in cultured lymphocytes and a lymphoblastoid cell line (Stillman et al., 1997; Zhang et al., 1998b). Other studies reported increased aneusomy of chromosomes 7 and 8 in cord blood CD34+ cells and loss of chromosome 7 together with deletion of 5q31 in CD34+CD19− bone marrow cells (Smith et al., 2000; Stillman et al., 2000). However, these experiments relied on FISH probes targeting a limited number of chromosomes and therefore could not establish the genome wide pattern of alterations. The relatively small numbers of nuclei analyzed and the potential for signal overlap or other technical artifacts also complicate the interpretation of low frequency chromosome loss.

The reported effects of HQ vary across studies. HQ and benzenetriol were found to induce aneuploidy involving nine chromosomes, suggesting a broader effect than selective targeting of chromosome 7 (Zhang et al., 2005). Other work emphasized chromosome breakage and hyperdiploidy (Eastmond et al., 1994). A micronucleus assay across eight donors detected no induction at HQ concentrations up to 200 µM, despite a positive response to a spindle poison control (Doepker et al., 2000). HQ also did not induce centrosome amplification in a study comparing it with formaldehyde (Ji et al., 2015). Although the absence of centrosome amplification does not exclude chromosome missegregation through other mechanisms, these findings leave the effects of HQ on chromosome segregation unresolved.

An alternative explanation is that HQ favors the survival or expansion of cells carrying chromosome loss. This possibility was raised by the original investigators, who proposed that the enrichment of chromosome 5 and chromosome 7 loss in benzene associated MDS and AML could reflect selective survival following HQ exposure (Stillman et al., 1997). Monosomy 7 clones can expand under hematopoietic stress, including stimulation with granulocyte colony stimulating factor (G-CSF) (Sloand et al., 2006). In germline SAMD9 and SAMD9L syndromes, these clones may persist, expand, or spontaneously regress, further suggesting that their fitness depends on the cellular and marrow environment (Wong et al., 2018; Sahoo et al., 2021).

Here, we asked whether HQ increases whole chromosome aneuploidy, particularly monosomy 7, in primary human CD34+ HSPCs at concentrations compatible with cell survival. We used KaryoTap to measure chromosome copy number across all 22 autosomes in thousands of single cells and performed chromosome 7 centromeric FISH on the same cultures as an orthogonal readout (Mays et al., 2024). Reversine, an inhibitor of the mitotic kinase MPS1, served as a positive control for chromosome missegregation in the cord blood experiment (Santaguida et al., 2010). The cord blood preparation contained two pooled donors, which we separated computationally using germline allele frequencies. This allowed us to assess donor specific responses within the same culture, although the donors did not represent independent experimental replicates. We then extended the analysis to adult mobilized peripheral blood CD34+ cells at two HQ concentrations and two timepoints. This approach allowed us to test for treatment associated changes in aneuploidy while accounting for baseline copy number calls and the response to a known inducer of chromosome missegregation.

## Results

### Hydroquinone toxicity defines the concentration range for single cell analysis

To test whether HQ induces chromosome missegregation, we first identified concentrations that would allow sufficient survival of CD34+ HSPCs for subsequent single cell analysis. Excessive cell death could eliminate cells with induced aneuploidy and limit our ability to detect these events. We cultured primary human CD34+ cells in serum free expansion medium and exposed them to 5 to 320 µM HQ for four days, assessing cell viability by propidium iodide (PI) exclusion. Cell death increased with HQ concentration, from 13.5% in the DMSO control to 14.9%, 19.7%, and 36.9% at 5, 10, and 20 µM, respectively. At 40 µM, cell death increased sharply to 92.4%, reaching 95 to 98% at concentrations of 80 µM and above (Fig. 1B; means of three replicates). We therefore selected 20 µM as the main working concentration, as this was the highest concentration tested that preserved sufficient viable cells for single cell analysis.

**Figure 1.**
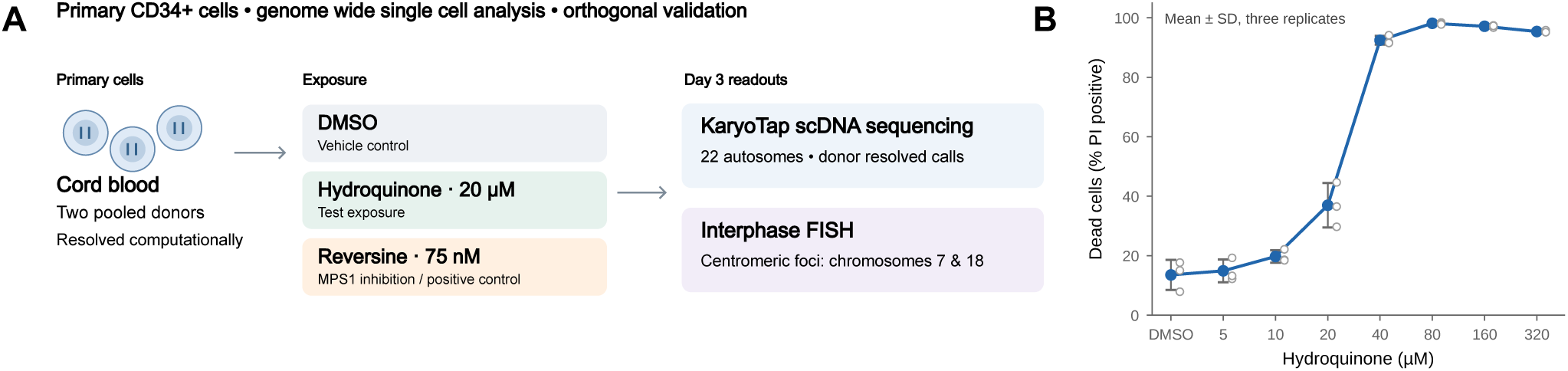
Experimental design and hydroquinone dose response. (A) Cord blood CD34+ HSPCs pooled from two donors were expanded in serum free medium and treated with DMSO, 20 µM HQ, or 75 nM reversine. Cultures were harvested at day 3 for KaryoTap analysis across the 22 autosomes and interphase FISH using centromeric probes for chromosomes 7 and 18. (B) Cell death after four days of exposure to DMSO or 5, 10, 20, 40, 80, 160, or 320 µM HQ, measured as the percentage of PI positive cells. Closed symbols and error bars indicate the mean and standard deviation of three replicates; open symbols show individual replicates.

Although HQ concentrations in the blood of benzene exposed workers are lower, approximately 0.2 to 1.1 µM, HQ can accumulate in the bone marrow. Marrow concentrations of 10 to 60 µM have been reported in rats exposed to high airborne benzene levels, providing the basis for estimates of approximately 10 to 20 µM in the marrow of highly exposed individuals (Ji et al., 2010). Our working concentration of 20 µM is at the upper end of this estimated range. It is below the 25 to 50 µM used in earlier in vitro studies reporting induction of monosomy 7; however, the marked toxicity observed at higher concentrations in our cultures limited our ability to examine their effects by single cell analysis.

### Reversine induces chromosome gains and losses in cord blood HSPCs

We treated cord blood CD34+ cells pooled from two donors with DMSO, 20 µM HQ, or 75 nM reversine and harvested the cultures at day 3 for KaryoTap and interphase FISH (Fig. 1A). Reversine was applied for approximately 39 hours, corresponding to the estimated doubling time of these cells, to allow cells to pass through mitosis during MPS1 inhibition. We selected 75 nM from a dose test spanning 50 to 300 nM. Concentrations of 150 nM and above caused visible stress and depletion, whereas 75 nM was tolerated.

The two donors separated clearly by their single cell allele frequency profiles in each condition and were analyzed independently (Fig. 2A). After demultiplexing and quality filtering, we retained 2,368 and 2,347 cells in DMSO, 3,011 and 3,332 cells in HQ, and 2,968 and 3,372 cells in reversine for donors 1 and 2, respectively.

**Figure 2.**
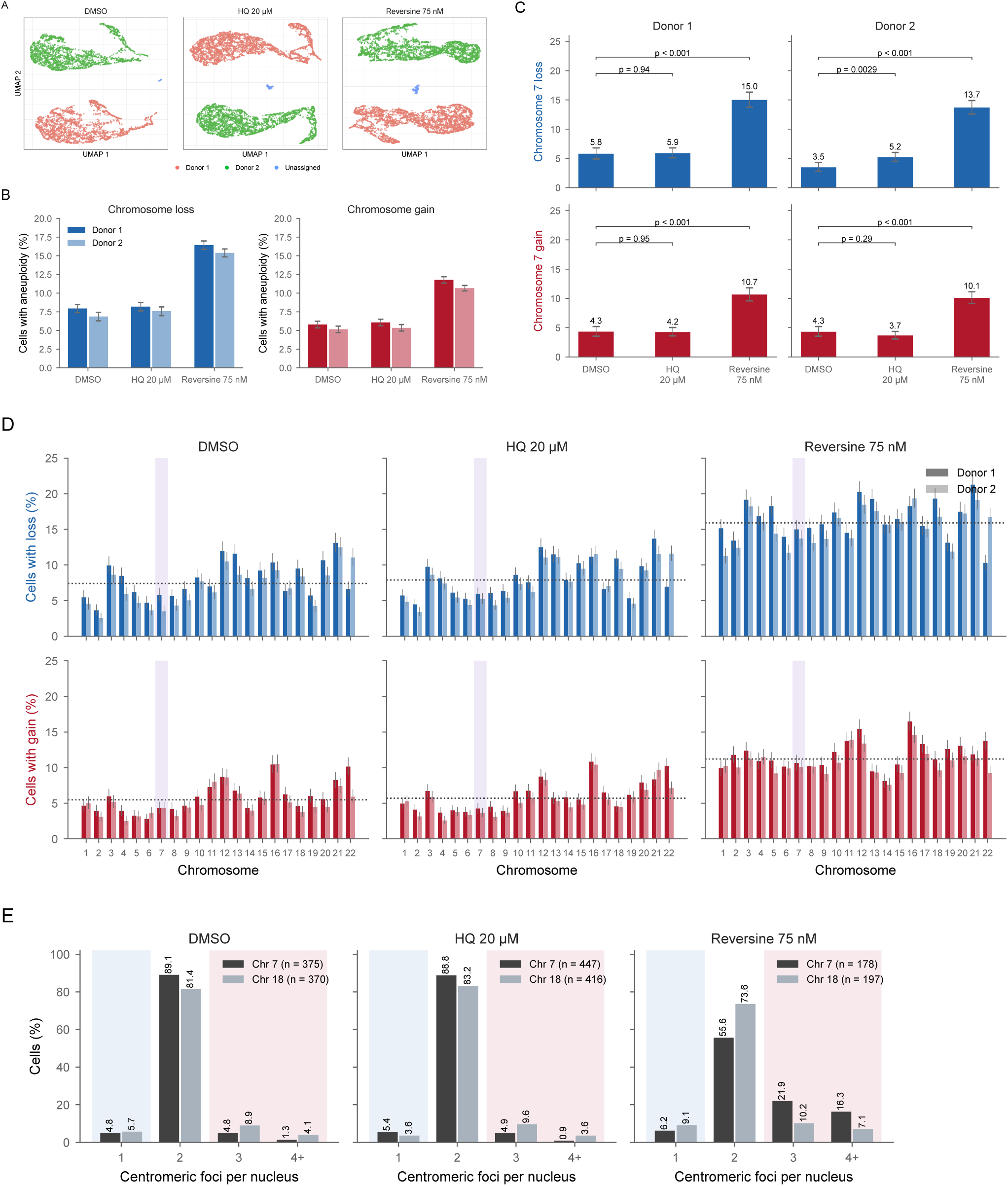
Reversine increases chromosome gains and losses in cord blood HSPCs at day 3. (A) UMAP projections of single cell allele frequency profiles for each condition, colored by donor assignment. The two pooled donors separate clearly, with a small number of cells remaining unassigned. (B) Frequency of chromosome loss (blue) and gain (red), averaged across the 22 autosomes, shown by condition and donor. Error bars indicate 95% confidence intervals. (C) Percentage of cells with chromosome 7 loss (blue) or gain (red), by condition and donor. Brackets show Fisher’s exact tests on cell counts with Benjamini Hochberg adjustment across 12 comparisons; the eight comparisons involving the vehicle control are displayed. Error bars indicate exact binomial 95% confidence intervals. (D) Percentage of cells with loss (upper row) or gain (lower row) of each autosome, by condition and donor. Whiskers indicate exact binomial 95% confidence intervals, and dotted lines show the mean across chromosomes within each panel. Chromosome 7 is highlighted. (E) Manual quantification of chromosome 7 and chromosome 18 centromeric FISH foci at day 3. Bars are colored by probe. Percentages are shown above the bars, and the number of nuclei scored for each probe is given in the key. Pale blue shading marks the one focus category; pale red shading marks the three focus and four or more focus categories.

Reversine increased chromosome gains and losses in both donors. Averaged across the 22 autosomes, the frequency of chromosome loss increased from 7.9% to 16.4% in donor 1 and from 6.9% to 15.4% in donor 2. The corresponding frequencies of chromosome gain increased from 5.8% to 11.8% and from 5.2% to 10.7% (Fig. 2B). The mean number of chromosomes lost per cell increased by 1.87 (95% CI, 1.67 to 2.06) in donor 1 and 1.88 (95% CI, 1.70 to 2.05) in donor 2. The mean number gained increased by 1.31 (95% CI, 1.22 to 1.41) and 1.21 (95% CI, 1.12 to 1.30), respectively.

The magnitude of the reversine response was similar between donors for both losses and gains (difference in differences p = 0.95 and p = 0.14, respectively). This agreement supports consistency across donor backgrounds, but the shared culture, treatment, and library preparation preclude treating the donors as independent experimental replicates. PI based DNA content analysis also showed a broader distribution after reversine treatment, with increasing disruption at higher concentrations in the dose test.

### Hydroquinone does not produce a comparable increase in whole chromosome aneuploidy

HQ treated cells showed copy number profiles similar to those of the DMSO controls. Averaged across the autosomes, chromosome loss frequencies were 8.2% and 7.6% after HQ, compared with 7.9% and 6.9% in DMSO for donors 1 and 2. Chromosome gain frequencies were 6.1% and 5.4% after HQ, compared with 5.8% and 5.2% in DMSO (Fig. 2B). Chromosome 5, another chromosome frequently examined in the benzene literature, showed a similar pattern. Loss frequencies were 6.1% and 5.4% after HQ, 6.2% and 4.7% after DMSO, and 18.3% and 14.4% after reversine in the two donors.

The chromosome specific profiles were also similar between HQ and DMSO across both donors, whereas reversine increased gains and losses broadly across the autosomes (Fig. 2D). Per chromosome values are provided in Supplementary Table S1 and shown as heatmaps in Supplementary Figure S1.

At harvest, viability was 88% in both HQ and DMSO cultures. Cell concentrations were 4.8 million cells/mL after HQ and 6.1 million cells/mL after DMSO. Reversine treated cultures had lower viability and cell concentrations, at 74% and 1.3 million cells/mL, respectively, despite showing a clear increase in aneuploidy. Thus, the absence of a comparable HQ response was not accompanied by a reduction in overall viability at harvest. These measurements do not exclude earlier or genotype specific differences in survival, but they argue against widespread cell death as the sole explanation for the result.

### Sequencing and FISH show no reproducible increase in chromosome 7 loss after hydroquinone

We next examined chromosome 7 specifically. Reversine increased chromosome 7 loss from 5.8% to 15.0% in donor 1 and from 3.5% to 13.7% in donor 2. Chromosome 7 gain increased from 4.3% to 10.7% and from 4.3% to 10.1%, respectively. All four comparisons were significant (p < 0.001, Fisher’s exact test with Benjamini Hochberg adjustment across 12 comparisons; Fig. 2C).

In donor 1, HQ had little effect on chromosome 7 copy number: loss frequencies were 5.9% after HQ and 5.8% after DMSO, and gain frequencies were 4.2% and 4.3%, respectively. In donor 2, chromosome 7 loss was modestly higher after HQ, at 5.2% compared with 3.5% in DMSO. This increase was not reproduced in donor 1 or in the adult donor at either concentration or timepoint. It was also smaller than the reversine response and was not accompanied by increased chromosome 7 gain, which decreased from 4.3% to 3.7%.

Reversine increased losses and gains in this donor by 10.2 and 5.8 percentage points, respectively. The absence of a parallel increase in gains does not by itself exclude missegregation, but the lack of replication limits the interpretation of the donor 2 result. In donor 1, the estimated upper bound on the HQ associated increase in chromosome 7 loss was approximately 1.4 percentage points, compared with an increase of approximately 9 to 10 percentage points after reversine.

We performed interphase FISH on the same cultures at day 3 using centromeric probes for chromosome 7 and chromosome 18, the latter as an internal control. Foci were counted manually in three fields per condition, comprising 178 to 447 cells per condition. In DMSO treated cells, 89.1% of nuclei had two chromosome 7 foci, 4.8% had one, and 6.1% had three or more. HQ treated cells showed a similar distribution: 88.8% had two foci, 5.4% had one, and 5.8% had three or more. After reversine, the proportion with two foci decreased to 55.6%, while 21.9% had three foci and 16.3% had four or more (Fig. 2E). The chromosome 18 probe also showed a shift only after reversine, consistent with a broad effect on chromosome segregation.

The FISH response to reversine was weighted toward increased foci counts, whereas KaryoTap detected more losses than gains. Differences in signal resolution, manual scoring, and the readouts themselves may contribute to this discrepancy. We therefore use FISH as qualitative support for the treatment response rather than as an independent quantitative estimate of chromosome gains and losses. Foci counts are provided in Supplementary Table S4.

Representative images from a separate day 10 imaging run are shown in Supplementary Figure S2.

### Adult mobilized HSPCs show no consistent hydroquinone response at either timepoint

We extended the analysis to adult mobilized peripheral blood CD34+ cells to test whether the result depended on the developmental source of the cells and to examine a later timepoint. The day 10 KaryoTap run from the cord blood experiment yielded fewer than 25 cells per condition, consistent with a technical failure during droplet encapsulation, and was excluded from interpretation.

CD34+ cells from an independent adult donor were treated with DMSO, 5 µM HQ, or 20 µM HQ for 72 hours. We analyzed samples designated day 3 and day 10 by KaryoTap. At day 3, we retained 5,529 cells: 1,951 in DMSO, 1,736 at 5 µM HQ, and 1,842 at 20 µM HQ. At day 10, we retained 4,848 cells: 1,469, 1,764, and 1,615, respectively. At day 3, chromosome 7 loss occurred in 1.6% of cells in each condition. Chromosome 7 gain frequencies were 4.3%, 4.0%, and 4.0% in DMSO, 5 µM HQ, and 20 µM HQ, respectively.

Frequencies averaged across the autosomes were also similar between conditions (Fig. 3; Supplementary Table S2). At day 10, chromosome 7 loss frequencies were 1.6%, 1.2%, and 2.0%, respectively. Neither chromosome 7 nor the autosomes as a whole showed a consistent dose dependent response at either timepoint. Together with the cord blood data, these results show no reproducible increase in whole chromosome aneuploidy or monosomy 7 across the three donors and two sources of CD34+ cells examined.

**Figure 3.**
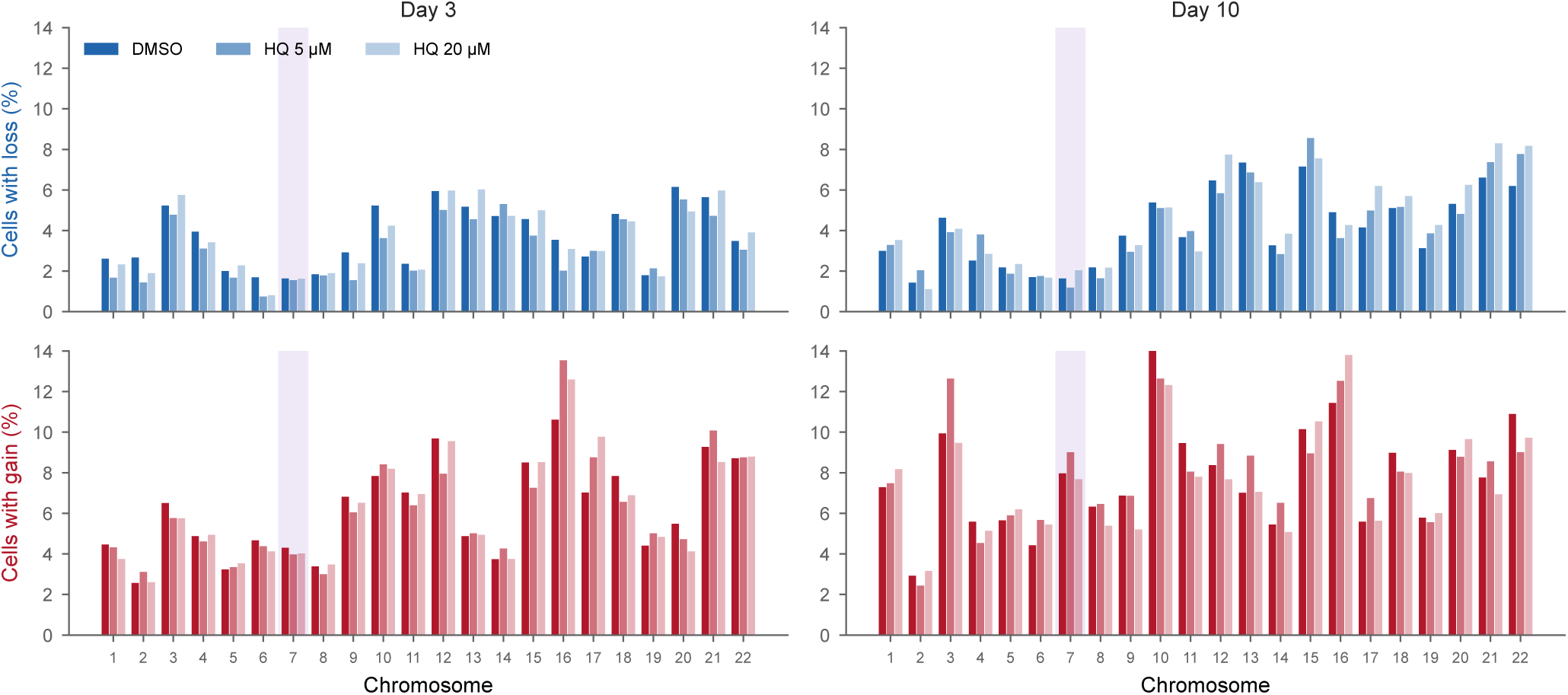
Hydroquinone does not consistently increase aneuploidy in adult mobilized HSPCs. Percentage of cells with loss (blue) or gain (red) of each autosome in adult mobilized peripheral blood CD34+ cells treated with DMSO, 5 µM HQ, or 20 µM HQ. Results are shown at day 3 and day 10, with chromosome 7 highlighted.

### HSPCs show a greater increase in chromosome losses after reversine than epithelial cells or melanocytes

We compared the reversine response in HSPCs with KaryoTap datasets generated in our laboratory from human mammary epithelial cells (hMECs) and melanocytes (hMELs) treated with the same 75 nM reversine regimen. Reversine increased the mean number of chromosomes lost per cell by 1.87 and 1.88 in the two HSPC donors, compared with 0.73 (95% CI, 0.63 to 0.84) in hMECs and 0.87 (95% CI, 0.80 to 0.95) in hMELs. The increase in HSPCs exceeded that in hMECs and hMELs by approximately 1.13 and 0.99 losses per cell, respectively (p < 0.001 for both comparisons; Fig. 4; Supplementary Table S3).

**Figure 4.**
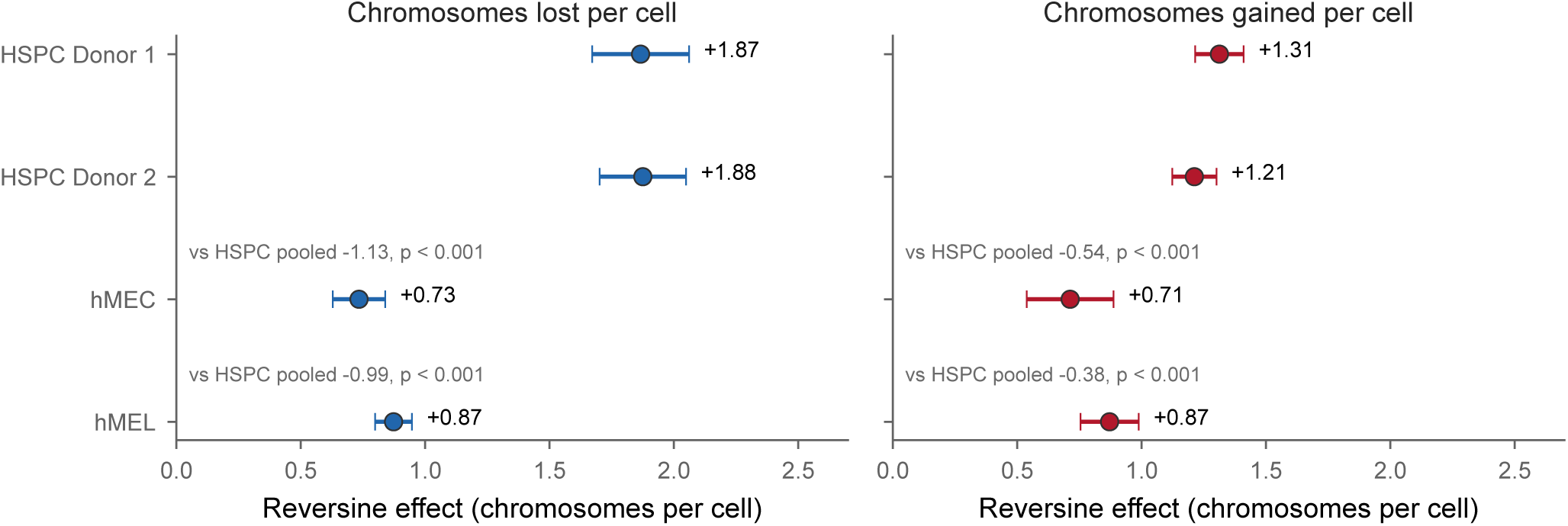
HSPCs show a greater increase in chromosome losses after reversine than epithelial cells or melanocytes. Change in the mean number of chromosomes lost (blue) or gained (red) per cell after reversine, relative to DMSO, in the two HSPC donors, human mammary epithelial cells (hMECs), and melanocytes (hMELs). Error bars indicate 95% confidence intervals. Contrasts between each comparison cell type and pooled HSPCs are also shown.

Differences in the increase in chromosome gains were smaller, at 0.54 and 0.38 chromosomes per cell relative to hMECs and hMELs. The cell type by treatment interaction terms differed in direction between losses and gains and were significant in three of four comparisons; the loss interaction comparing HSPCs with hMECs did not reach significance (Supplementary Table S3).

These results suggest that the response to reversine differs by cell type, with a greater accumulation of chromosome losses in HSPCs. This could reflect differences in missegregation, survival, or subsequent proliferation and does not establish greater tolerance of chromosome loss on its own. The comparison demonstrates that KaryoTap can resolve cell type associated differences in a strong treatment response. Sensitivity to small changes above baseline must instead be assessed from the uncertainty around the HQ comparisons.

## Discussion

Our results do not support a substantial, reproducible increase in whole chromosome aneuploidy following HQ exposure in primary human CD34+ HSPCs under the conditions tested. Across three donors, two sources of CD34+ cells, and more than 20,000 single cells, HQ did not consistently increase chromosome 7 loss or whole chromosome gains and losses more broadly. The adult experiment extended this result to two HQ concentrations and two timepoints. A modest increase in chromosome 7 loss was observed in one cord blood donor, but it was not reproduced in the other donors. In the cord blood experiment, reversine produced a clear increase in aneuploidy in the same cultures, confirming that the assay could detect a strong response to chromosome missegregation. This positive control does not exclude a smaller HQ effect below the sensitivity of the experiment.

Several factors may contribute to the differences between our findings and earlier reports. Previous studies primarily used FISH to examine selected chromosomes in a few hundred to a few thousand nuclei. Single signal nuclei can reflect true chromosome loss, but signal overlap, incomplete hybridization, and sample preparation artifacts can also affect these measurements. These limitations do not invalidate the earlier findings, but they make orthogonal, genome wide measurements valuable. Differences in cell source, culture conditions, concentration, and exposure duration may also contribute to the discrepancy.

Studies examining more chromosomes have generally reported broader patterns of aneuploidy rather than a selective effect on chromosome 7. In vitro, HQ and benzenetriol affected nine chromosomes (Zhang et al., 2005). In benzene exposed workers, an eight color FISH survey found dose dependent increases in monosomy of chromosomes 5, 6, 7, 10, 16, and 19 and trisomy of chromosomes 5, 6, 7, 8, 10, 14, 16, 21, and 22 (Zhang et al., 2011). Other assays have also produced mixed results: a study across eight donors found no micronucleus induction at HQ concentrations up to 200 µM, and HQ did not induce centrosome amplification in lymphoblastoid cells (Doepker et al., 2000; Ji et al., 2015). These findings do not exclude all possible effects on chromosome segregation, but they support a fresh examination of how HQ influences chromosome copy number in different experimental settings.

Our findings are consistent with a model in which benzene exposure favors the expansion of preexisting monosomy 7 cells. Under this model, HQ or other consequences of benzene exposure alter the marrow environment so that cells carrying chromosome 7 loss gain a relative fitness advantage. This possibility was proposed by the investigators who first reported HQ associated chromosome loss in a lymphoblastoid cell line (Stillman et al., 1997). It is also consistent with evidence that monosomy 7 clones can expand under hematopoietic stress, including G-CSF stimulation, and can follow different trajectories in SAMD9 and SAMD9L syndromes (Sloand et al., 2006; Wong et al., 2018; Sahoo et al., 2021). Our experiments did not directly test selection and cannot establish that it explains the association between benzene exposure and monosomy 7. They provide a rationale for testing that model directly.

### Limitations of this study

The highest HQ concentration we analyzed by KaryoTap was 20 µM, below the 25 to 50 µM used in earlier studies reporting monosomy 7 induction. We therefore cannot exclude an effect at higher concentrations. In our dose response experiment, 40 µM caused extensive cell death, limiting the number of surviving cells available for analysis. Although 20 µM is near the upper end of published estimates for marrow exposure in highly exposed individuals, those estimates are uncertain and do not reproduce the complexity of exposure in vivo. The cord blood experiment included one HQ concentration, while the adult experiment included two. Neither was designed to exclude very small changes above baseline.

The modest increase in chromosome 7 loss in cord blood donor 2 warrants follow up in independently cultured donors. Its absence in donor 1 and the adult donor limits its generalizability, but neither the smaller magnitude relative to reversine nor the absence of increased chromosome 7 gain is sufficient to dismiss it. The two cord blood donors shared culture, exposure, and library preparation, and therefore provide information about donor specific responses within one experiment rather than independent experimental replication. The adult validation used a single donor and did not include a concurrent reversine control.

The present analysis was restricted to whole chromosome copy number. We did not examine del(7q), del(5q), or other arm level alterations, although these have been reported after benzene or HQ exposure. KaryoTap supports arm level analysis, which would be a useful extension of this study. Copy number calling also has technical limitations, including probe dropout and differences in confidence between loss and gain calls. We therefore emphasize comparisons with concurrent vehicle controls rather than absolute aneuploidy frequencies or differences in baseline frequencies between experiments.

We did not measure a direct marker of HQ exposure or activity in the cultures used for sequencing. HQ oxidizes readily in bicarbonate buffered medium, so the effective concentration during the 72 hour exposure may have differed from the nominal concentration despite supplementation at medium changes. The dose dependent toxicity observed with the same preparation supports compound activity but does not establish the exposure achieved in each sequencing sample. A damage response marker such as γH2AX, measured in the same cultures, could provide additional evidence of biological activity; direct measurement of HQ would be needed to establish the concentration over time.

Technical failure prevented interpretation of the cord blood day 10 KaryoTap data, so evidence at the later timepoint comes only from the adult experiment. FISH provided an orthogonal readout but was scored manually in a limited number of fields and showed a different balance of gains and losses after reversine. We therefore interpret it as qualitative support rather than an independent quantitative estimate.

Finally, these experiments used defined culture conditions without marrow stroma or the inflammatory signals present in vivo. They cannot reproduce the full effects of benzene exposure, including the contributions of other metabolites. Although the design reduces some sources of environmental selection, differential survival and proliferation can still occur in culture. Endpoint copy number measurements therefore reflect both the generation and persistence of aneuploid cells and cannot fully separate induction from selection.

The greater accumulation of chromosome losses in HSPCs after reversine raises a separate question about how cell type influences the response to aneuploidy. Distinguishing differences in chromosome missegregation from differences in survival or proliferation will require direct measurements of these processes. This may be particularly relevant to understanding why chromosome losses recur in myeloid disease.

A direct test of the selection model would compare the fitness of monosomy 7 and euploid cells under HQ exposure. KaryoCreate provides a means to engineer chromosome specific aneuploidy in human cells and could support this approach, subject to validation in the relevant HSPC system (Bosco et al., 2023). Monosomy 7 cells could then be mixed with euploid counterparts and followed over time by single cell sequencing or a validated lineage tracking method, under HQ exposure and defined inflammatory conditions. Combining these competition experiments with direct measurements of chromosome missegregation would help distinguish effects on lesion formation from effects on clonal expansion. The same strategy could be applied to other exposures implicated in myeloid disease, including alkylating agents and chronic inflammatory stimuli.

## Methods

### Cells and culture

Human CD34+ cells were obtained from STEMCELL Technologies. The cord blood preparation (catalog 70008, lot 2505510003) contained cells pooled from two donors, as confirmed by the supplier. These donors were separated computationally and analyzed individually, but shared culture and experimental processing. Adult mobilized peripheral blood CD34+ cells were obtained from a single 43 year old male donor. Cells were thawed into StemSpan serum free expansion medium and allowed to recover and expand for two to three days before exposure. Unless otherwise specified, day numbering is relative to the start of HQ treatment: day 3 corresponds to the end of the 72 hour exposure, and day 10 to one additional week of culture.

### Hydroquinone exposure

A 20 mM HQ stock was prepared in DMSO, aliquoted, and stored at −20 °C protected from light. Working solutions of 0.5 mM and 2 mM HQ were prepared fresh in PBS on the day of exposure, together with a DMSO vehicle solution. Working solutions were added at 1:100 to achieve final HQ concentrations of 5 µM and 20 µM. The final DMSO concentration was reported as 0.1% in all conditions, including the vehicle control.

Cells were pelleted at 320 × g for 5 minutes and resuspended in HQ containing medium for 72 hours. HQ was replenished at each medium change. In the adult experiment, exposure began three days after thawing, and cells were subsequently returned to standard medium. Samples were designated day 3 and day 10 for analysis.

### Reversine exposure and dose selection

A 10 mM reversine stock was diluted to 100 µM in PBS and added to cultures at a final concentration of 75 nM. Cells were exposed for approximately 39 hours, corresponding to their estimated doubling time, to allow mitotic progression during MPS1 inhibition. Cells were then pelleted and transferred to fresh StemSpan medium. The working concentration was selected from a parallel dose test using cells from the same thaw and concentrations of 50, 75, 150, and 300 nM. Cultures treated with 150 nM or more showed marked depletion and abundant cellular debris, whereas 75 nM was tolerated.

### Viability and DNA content measurements

For the HQ dose response experiment, cells were exposed for four days to DMSO or 5, 10, 20, 40, 80, 160, or 320 µM HQ, with three replicates per concentration. Viability was measured by PI exclusion. For the reversine dose test, cells were fixed in 100% ethanol, treated with RNase A, stained with PI for one hour, and analyzed by flow cytometry. DNA content distributions were compared across concentrations.

### Single cell DNA sequencing

Single cell copy number was measured using the Mission Bio Tapestri DNA and Protein platform with the KaryoTap panel. The three conditions in each experiment were multiplexed by antibody hashing. Cells were blocked on ice for 15 minutes with blocking buffer and Human TruStain FcX, then stained with one TotalSeq-D hashing antibody per condition at 0.25 µg per clone for 30 minutes on ice in the dark. Cells were washed twice in cell staining buffer, passed through a 40 µm strainer, counted, and pooled at a 1:1:1 ratio. Approximately 83,000 cells per condition were pooled at a final concentration of 4,000 to 5,000 cells/µL. Cells were encapsulated and barcoded according to the Tapestri protein and DNA protocol. DNA and protein libraries were quantified by Qubit and TapeStation and pooled for sequencing at 5 nM and 0.15 nM, respectively.

### Copy number calling and donor demultiplexing

Reads were processed using the Mission Bio pipeline, and cells were assigned to conditions using their hashing antibodies. In the cord blood experiment, cells were further assigned to donors using allele frequencies at germline variant sites. Donors were analyzed separately. Cells were filtered by total reads per cell, and doublets were excluded.

Copy number was normalized using the KaryoTap workflow, which produced a narrower copy number distribution in DMSO treated cells than standard pipeline normalization. A Gaussian mixture model was then applied to smoothed per chromosome copy number values. A chromosome was classified as lost or gained when its called copy number was below or above two, respectively.

Posterior probabilities from the mixture model were not fully calibrated. Copy number distributions widened at higher copy number states, and probe dropout produced low values in individual cells, resulting in systematically higher nominal confidence for loss calls than for gain calls. We therefore compared each treatment with its concurrent DMSO control and did not interpret absolute frequencies as direct estimates of biological aneuploidy. Baseline frequencies also differed between experiments and were not compared directly.

### Fluorescence in situ hybridization

Cells were attached to poly-L-lysine coated slides, fixed in 4% paraformaldehyde, washed in PBS, and stored at 4 °C. Hybridization used a centromeric probe for chromosome 7 and a commercial chromosome 18 centromeric probe as an internal control (Empire Genomics, Chr18-10-GR). Probes were used at 6 pmol per 30 µL of hybridization buffer, with 25 µL applied per spot. Slides were denatured for 5 minutes and hybridized overnight at 42 °C in a humidified chamber. Slides were then washed, mounted, and allowed to cure before imaging. Foci were counted manually in three fields per condition, comprising 178 to 447 cells per condition. Each nucleus was classified as having one, two, three, or four or more foci for each probe.

### Statistics

Differences in the proportion of cells carrying a given event were tested using Fisher’s exact test on cell counts, with Benjamini Hochberg adjustment across the full set of comparisons within each figure. Proportions are reported with binomial 95% confidence intervals. Differences in the number of chromosomes gained or lost per cell were tested using the Wilcoxon rank sum test on individual cells within each system. Reversine effects are reported as changes in the mean number of altered chromosomes per cell, with 95% confidence intervals. Treatment responses were compared between cell systems and between the two cord blood donors using difference in differences analyses.

Tests based on individual cells measure differences within the sampled cultures and do not establish reproducibility across independent cultures or donors. We therefore interpret these results alongside agreement between donors and experiments rather than relying on nominal significance alone.

## Acknowledgements

We thank all members of the Davoli lab for helpful comments and discussions. We also thank William Carroll, Christopher Park, Iannis Aifantis, and David Beck, members of M.T.’s thesis committee, for their advice throughout this work. This study was supported by the National Institutes of Health through R37 CA248631 from the National Cancer Institute and R01 HG012590 from the National Human Genome Research Institute, and by the Pershing Square Sohn Prize for Young Investigators in Cancer Research. We apologize to colleagues whose work we could not cite because of space constraints.

## Author contributions

M.T. designed and performed experiments and analyzed the data. S.A. made essential contributions to the HQ exposure and FISH experiments. R.K. contributed to the FISH and sequencing experiments. F.A. and M.K. contributed to single cell sequencing and computational analysis. T.D. conceived and supervised the study. M.T., T.D., and S.A. wrote the manuscript with input from all authors.

## Data and code availability

The single cell DNA sequencing data and per chromosome copy number calls summarized in Supplementary Tables S1 and S2 will be deposited in a public repository upon publication. The analysis code will also be made publicly available. Additional information needed to reanalyze the data is available from the lead contact upon request.

## Figure legends

Chromosome losses are shown in blue and gains in red throughout. Conditions or donors are distinguished by shading within each color. In Figure 2E, bars are colored by probe rather than event type.

**Supplementary Figure S1. Chromosome copy number profiles across conditions.** (A) Heatmaps showing the difference between the percentage of cells with a gain and the percentage with a loss for each chromosome. The upper heatmap shows the cord blood experiment at day 3 by condition and donor; the lower heatmap shows the adult mobilized experiment by concentration and timepoint. (B) Cord blood data from panel A shown separately as loss and gain frequencies. Chromosome 7 is outlined throughout. Each heatmap has its own color scale, so color intensity should not be compared across heatmaps. The cord blood experiment had more loss than gain calls at baseline, whereas the adult experiment had more gain than loss calls. Treatment effects should therefore be assessed by comparing rows within each heatmap. HQ and vehicle profiles are similar, while reversine produces a clear shift.

**Supplementary Figure S2. Representative chromosome 7 FISH images.** Maximum intensity projections of representative fields from CD34+ HSPC cultures imaged at day 10 after treatment with DMSO, 20 µM HQ, or 75 nM reversine. DAPI is shown in grey and the chromosome 7 centromeric probe in green. Boxed regions are enlarged below each field. These images were obtained in a separate day 10 imaging run in which only the chromosome 7 probe was acquired. They illustrate hybridization appearance and were not used for the day 3 quantification in Figure 2E, which included the chromosome 18 control probe.

## Supplementary table legends

**Table S1. Chromosome copy number in the cord blood experiment.** Number of cells analyzed and percentage with a loss, gain, or any non diploid call, by condition, donor, chromosome, and timepoint. Day 10 values are included for completeness but should not be interpreted because technical failure left fewer than 25 cells per condition.

**Table S2. Chromosome copy number in the adult mobilized experiment.** Per chromosome copy number summaries for DMSO, 5 µM HQ, and 20 µM HQ at day 3 and day 10.

**Table S3. Reversine responses across cell systems.** Changes in the mean number of chromosomes lost or gained per cell, with standard errors and confidence intervals; contrasts between each comparison system and pooled HSPCs; the contrast between HSPC donors; and cell type by treatment interaction terms.

**Table S4. Manual FISH counts.** Counts of chromosome 7 and chromosome 18 centromeric foci per nucleus at day 3, by condition.

