## Supplementary figures and images for "KaryoTap-based scDNA seq assessment of aneuploidy induction by hydroquinone in hematopoietic stem cells"

### Suppl Figures

#### Supplementary Figure S1

A

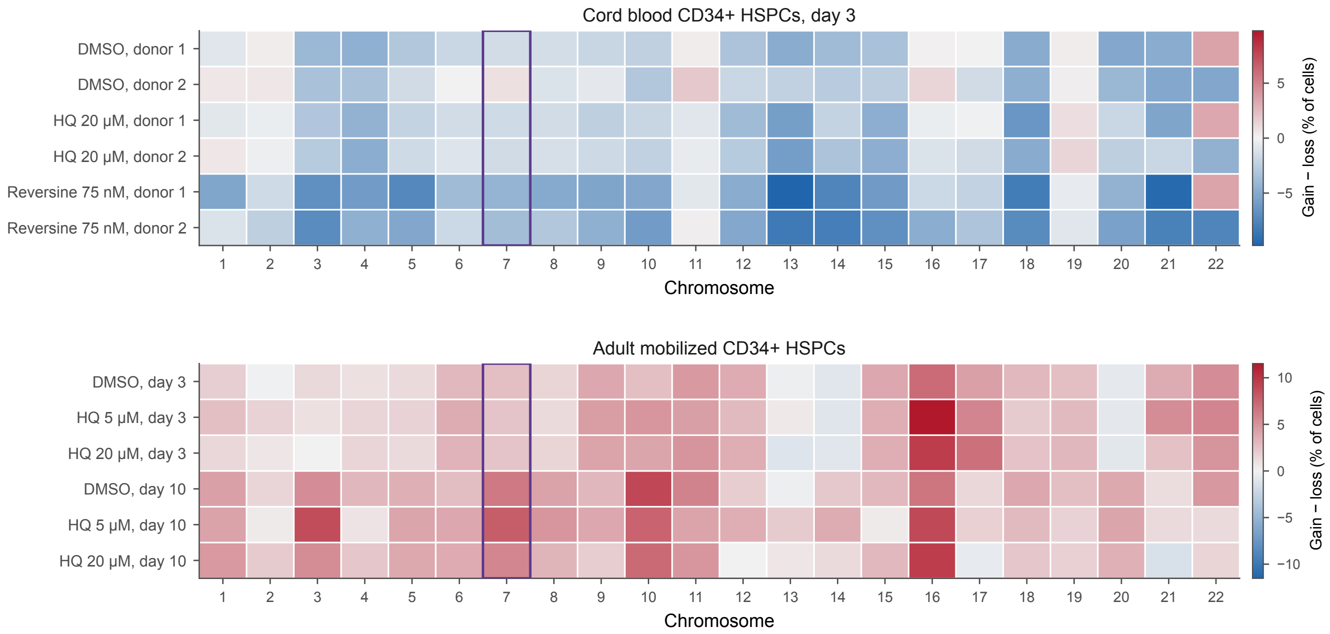


B

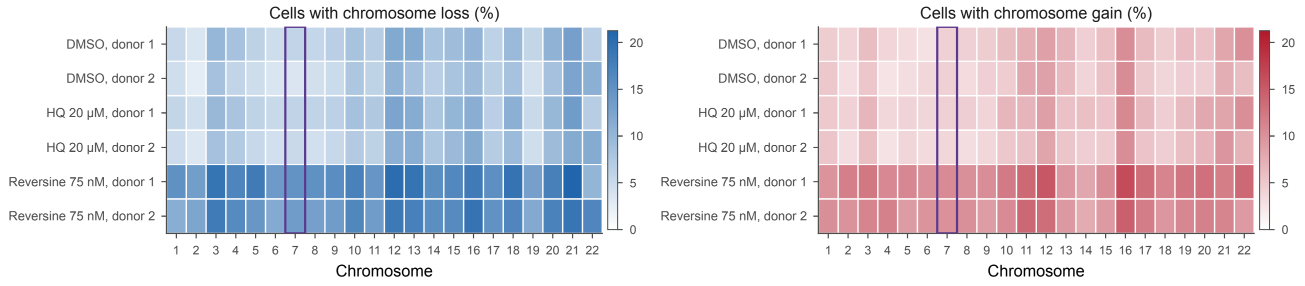

#### Supplementary Figure S2

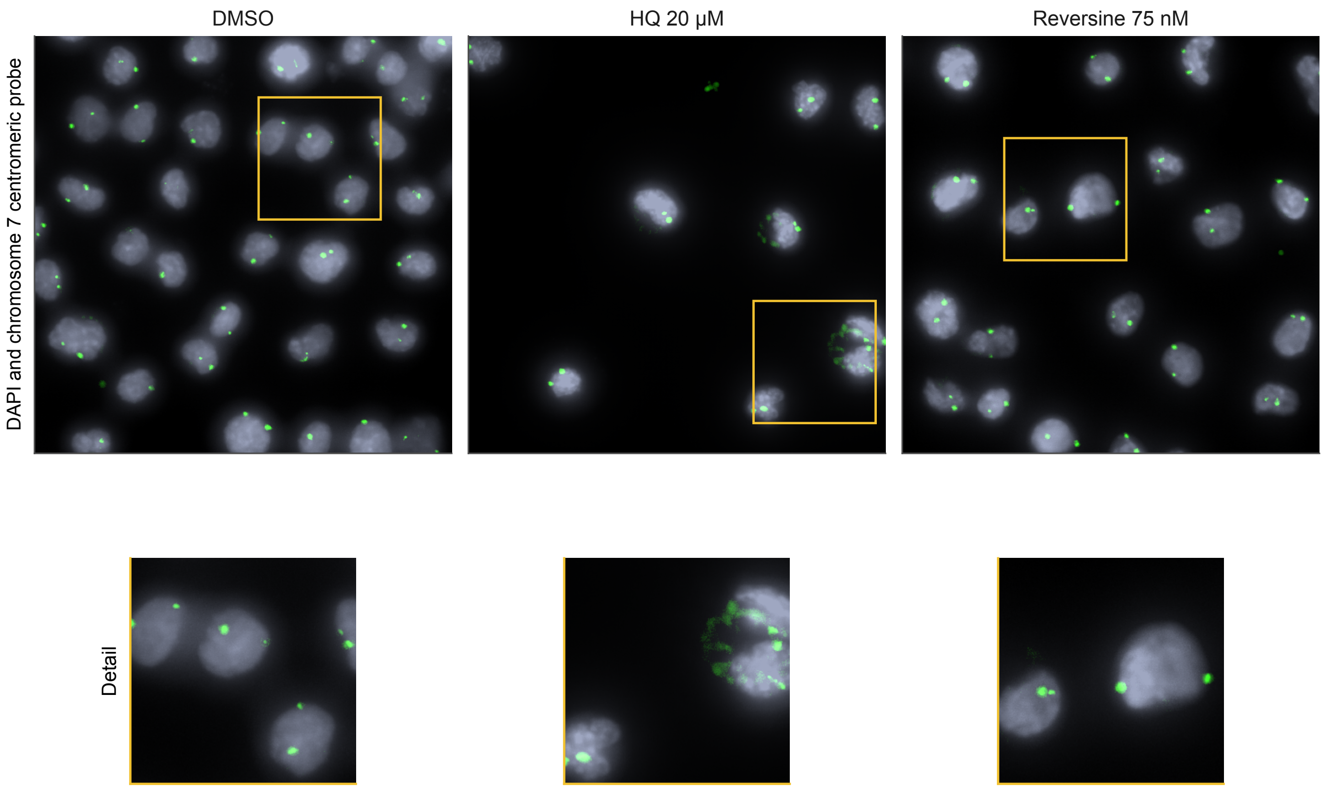
